# Hibernation improves neural performance during energy stress in regions across the central nervous system in the American bullfrog

**DOI:** 10.1101/2025.06.13.659541

**Authors:** Nikolaus Bueschke, Joseph M. Santin

## Abstract

Neuronal signaling requires high rates of ATP production via the oxidative metabolism of glucose. The American bullfrog is intriguing, as this species has typical brain energy requirements for an average vertebrate but modifies synaptic physiology and metabolism after hibernation to maintain function during hypoxia and ischemia. Given the importance of the respiratory system in restoring metabolic homeostasis during emergence from underwater hibernation, work to date has addressed this response in the brainstem respiratory network. Thus, metabolic plasticity has been interpreted as an adaptation used to restart respiratory motor behavior under hypoxic conditions during the transition from skin breathing to air breathing. It remains unclear whether these improvements are specific to the brainstem regions critical for breathing *versus* a global response within the central nervous system (CNS). To address this question, we recorded neural activity from the spinal cord, forebrain, and brainstem respiratory network *in vitro*. As expected, hypoxia disrupted the function of each network in control animals. After hibernation, each network improved its activity in hypoxia compared to controls. These results suggest that plasticity that improves neural function during energy stress following hibernation reflects a global response that may impact many behaviors controlled by the CNS and is not limited to regions involved in metabolic homeostasis.

## Introduction

Neural function is energetically costly. Energy deficits during hypoxia or ischemia rapidly lead to brain dysfunction and eventual neuronal cell death (Buck & Pamenter, 2018). While this holds for humans and laboratory mammals, several animals have evolved mechanisms to overcome low levels of O_2_ that occur as a part of their normal life histories (Larson et al., 2014). For example, freshwater turtles can survive anoxic environments during prolonged periods (Jackson & Ultsch, 2010) and respond by decreasing excitatory neurotransmitter release, increasing inhibition, leading to a controlled and reversible coma-like state with reduced neuronal function to preserve energy (Fernandes et al., 1997). Additionally, hibernating arctic ground squirrels suppress NDMA-glutamate receptor expression to compensate for anoxia-induced glutamate release to reduce function in events of limited energy supply (Jinka et al., 2012; Zhao et al., 2006). While spike and synaptic arrest are common occurrences that likely aid in neuronal protection in hypoxia, the strategies for tolerance in species can vary depending on life history (For review, see Larson et al., 2014).

Unlike many of these examples of naturally evolved hypoxia tolerance, amphibians hold a different position along the spectrum of hypoxia tolerance. These animals have typical brain energy requirements for an average vertebrate but dramatically improve neural performance under energy stress following hibernation (Bueschke, Amaral-Silva et al., 2021). Indeed, brainstem circuits of the American bullfrog are not tolerant to hypoxia or ischemia and fail within minutes of exposure through mechanisms consistent with metabolic failure rather than modulatory suppression of the neurons (Adams et al., 2021). However, after aquatic overwintering, these animals undergo metabolic changes that shift brainstem circuits to maintain function in anoxia and oxygen and glucose deprivation for over 2 hours. The mechanism for metabolic plasticity involves changes in the primary modes of ATP generation, where anaerobic glycolysis becomes sufficient to fuel brainstem motor networks without oxygen or glucose delivery, using only glycogen as a carbon source (Bueschke, Amaral-Silva et al., 2021). Metabolic adjustments appear to support synaptic activity, rather than action potential firing and subsequent ion regulation, as synaptic transmission is limiting to network function during hypoxia (Amaral-Silva & Santin, 2023). To improve ward off excitotoxic activity patterns in hypoxia, NMDA-glutamate receptors reduce Ca^2+^ permeability and increase desensitization while keeping overall expression levels the same, which maintains the normal role of NMDA receptors in motor output in well-oxygenated conditions, but reduces hyperexcitability during hypoxia (Bueschke, Amaral-Silva et al., 2024). Therefore, several metabolic and neurophysiological mechanisms improve the capacity of the brainstem to improve its function without oxygen and/or glucose delivery.

Why are these processes adaptive for the hibernating frog? When frogs are in the submerged hibernation environment, blood O_2_ levels are low (1-5 mmHg compared to 80-90 mmHg; Tattersall & Ultsch, 2008; Ultsch et al., 2004). Low O_2_ levels are not immediate concerns given the low rates of metabolism in the cold. However, limitations in O_2_ become more problematic during emergence as the functional demands of the nervous system rise and temperature increases (Amaral-Silva & Santin, 2024). Given the central position of the respiratory system in restoring whole body metabolic homeostasis, the primary focus of our previous work has been on the brainstem respiratory network, as this is a critical neural system that must restart on the background of hypoxia during emergence to restore blood gas homeostasis. However, the vertebrate nervous system has a wide array of differing responses to anoxia and ischemia, leaving to question whether these responses are specific to the brainstem or a global response of the central nervous system. For example, at least in mammals, brainstem and pontine neurons tend to survive ischemia better than neurons of the cortex, hippocampus, and striatum, which is believed to be induced in part by Na^+^/K^+^ ATPases in the hippocampus, making them more susceptible to failure in ischemic conditions (Brisson et al., 2014). Even within brain regions, differences in sensitivity to ischemic injury exist. For example, CA1 neurons found in dorsal hippocampal neurons suffer greater degrees of cell death compared to those in the ventral region, potentially due to differences in NMDA-glutamate receptor subtypes (Ashton et al., 1989; Hurley et al., 2022). Additionally, alpha motor neurons resist ischemia better than in sensory neurons since motor neurons have a slower depolarization and slower sodium channel inactivation (Hofmeijer et al., 2013). Because of these differences in sensitivity to energy stress across the central nervous system, it is important to investigate how the metabolic plasticity in the brainstem translates to other parts of the nervous system.

To build upon our work in the brainstem, we assessed the ability for the spinal cord and forebrain synapses to operate during energy stress before and after hibernation. Given the wealth of literature pointing to regional differences in sensitivity to hypoxia and ischemia, we hypothesized that energy stress in the forebrain and spinal cord would exhibit differing responses compared to the brainstem and that plasticity responses would differ as well. We focused on two additional regions within the central nervous system and how their response to energy stress changes following hibernation: the spinal cord and forebrain. In vitro, the spinal cord produces rhythmic activity when descending inputs from the brainstem are intact (Rauscent et al., 2006; Sutherland & Nunnemacher, 1974). Therefore, we tested tolerance to hypoxia through extracellular nerve root recordings in a preparation that includes both the brainstem and complete spinal cord. For forebrain function, we assessed synaptic transmission in neurons of the dorsal pallium, which is believed to be involved in associative learning in amphibians (Roth et al., 2003, 2007; Sotelo et al., 2024; Woych et al., 2022). We tested tolerance to energy stress by measuring changes in evoked synaptic response in the presence of OGD using whole-cell patch clamp electrophysiology.

## Methods

### Animal husbandry and ethical approval

The use of animals was approved by the Animal Care and Use Committee (ACUC) at the University of Missouri (Protocol #39,264). Female American bullfrogs (∼ 100 g, n=43 were shipped from Frog Farm (Twin Falls, ID, USA). Frogs were assigned randomly between control and hibernation groups and placed into tubs containing 20 gallons of dechlorinated water. Control groups were held at ambient room temperature (∼20°C) with access to a floating platform and fed pellets supplied by Phrog Farm weekly. Control frogs were given at least 1 week to acclimate following arrival before experimental use. Hibernation groups were exposed to temperatures that were gradually decreased from 20°C to 4°C over the course of 7 days using a walk-in temperature-controlled environmental chamber. Once at 4°C, floating platforms were removed and hibernating frogs were kept submerged by adding a plastic screen below the surface to promote cutaneous respiration and held at temperature for 30 days. In both groups, frogs were kept at 12:12 hour light/dark cycles and water was kept aerated by aquarium air pumps.

### Dissection and slice preparation

Frogs were anesthetized with isoflurane (∼1 ml/L) and rapidly decapitated after loss of toe-pinch response. Heads were then submerged in chilled artificial cerebral spinal fluid (aCSF; 104 mM NaCl, 4 mM KCl, 1.4 mM MgCl2, 7.5 mM D-glucose, 1 mM NaH2PO4, 40 mM NaHCO3, and 2.5 mM CaCl2) bubbled with 98.5% O_2_ and 1.5% CO_2_ resulting in a pH of 7.9 ± 0.1. This pH and CO_2_ concentration is normal for frogs at room temperature (Howell et al., 1970). For extracellular recordings of the respiratory motor network and the caudal spinal cord network, the entire brainstem and spinal cord was rapidly exposed to oxygenated aCSF and carefully excised from the cranium. For patch clamp recordings of the dorsal pallium, the forebrain was removed similarly to the spinal cord.. In both dissections, the dura membrane was removed. The forebrain was then transferred to a vibrating microtome (Campden Vibrating Microtome 7000smz, Campden Instruments; Lafayette, IN, USA), where transverse slices were made at 300 µm sections from rostral to caudal in oxygenated ice-cold aCSF. The brain was kept upright by gluing the ventral side to a block of agar. Slices were cut in half to isolate hemispheres and held at room temperature, oxygenated aCSF (22±1°C).

### Extracellular nerve rootlet recordings

Immediately following dissection, brainstem-spinal cord preparations were transferred to 6 mL petri dishes coated with Sylgard (Dow Inc. Midland, MI, USA) where they were pinned ventral side up and continuously superfused with room temperature aCSF oxygenated with the control gas mixture (98.5% O_2_, 1.5% CO_2_) using peristaltic pumps (Watson-Marlow, Falmouth, UK; Rainin Instrument Co. Inc). For nerve recordings, suction electrodes were made from borosilicate glass pulled from a horizontal puller (P-1000 Micropipette Puller, Sutter Instruments, Novato, CA, USA), carefully broken at the tip, and fire-polished to establish a tight seal for nerve root attachment. Rhythmic output from the respiratory motor network and the Spinal cord were recorded simultaneously. Respiratory output was recorded from trigeminal (CN V), vagus (CN X), and hypoglossal (CN XII) rootlets. Rhythmic spinal cord output was recorded through SN III, and SN VII-IX rootlets. These spinal nerves innervate the limbs of anuran amphibians (SN III to the forelimb, SN VII-IX to the hindlimb; Sutherland & Nunnemacher, 1974). Signals were amplified (x100), filtered (low-pass, 1000 Hz, high-pass, 10 Hz) using an amplifier (A-M Systems 1700 amplifier, Sequim, WA, USA), and digitized with a PowerLab 8/35 (ADInstruments, Dunedin, New Zealand). Signal was recorded at 1 k/sec on LabChart 8 software (ADInstuments, Dunedin, New Zealand) where the raw signal was rectified and integrated. Anoxia was induced by replacing oxygenated aCSF with aCSF bubbled with N_2_, replacing O_2_ (98.5% N_2_, 1.5% CO_2_).

### Patch clamp electrophysiology and stimulation protocols

Neurons in the dorsal pallium were approached with thin-walled borosilicate glass pipettes (3-5 MΩ). Whole-cell recordings were made from dorsal pallium neurons with an Axopatch 200B amplifier (Molecular Devices, San Jose, CA, USA). Pipettes were filled with intracellular solution (110 mM K-gluconate, 2 mM MgCl_2_, 10 mM HEPES, 1 mM Na_2_-ATP, 0.1 mM Na_2_-GTP, 2.5 mM EGTA, pH to ∼7.2 with KOH). Slices were perfused with aCSF and oxygenated with 98.5% O_2_ and 1.5% CO_2_. OGD was induced by perfusing in glucose-free aCSF, where glucose was replaced with equimolar sucrose bubbled with 98.5% N_2_ and 1.5% CO_2_. Preliminary experiments suggested perturbations for 15 minutes were optimal for maximizing throughput on continuously recorded neurons. Due to time constraints, we wanted to ensure the strongest perturbation to energy possible, OGD. Cells were exposed to OGD perfusion for 15 minutes before running OGD protocols. Neurons were held at the reversal potential of Cl^-^ (−92 mV) to isolate excitatory neurotransmission. Stimulating pulses at 0.5 Hz were applied with gradually increasing currents until recorded neurons displayed a consistent maximal evoked EPSC to ensure saturation (∼10-50 mA stimulus output). Upon saturated evoked current, neurons remained stimulated at 0.5 Hz for at least 60 seconds before data collection to ensure stability.

Bipolar tungsten stimulating electrodes (MicroProbes for Life Science, Gaithersburg, MD, USA) were placed on the distal edge of slices and placed on thalamic projections that innervate recorded neurons (Roth et al., 2003). Simulating current was driven by an isolated pulse stimulator (A-M Systems Model 2100, Sequim, WA). Pipettes and stimulating electrodes were driven with a micromanipulator (MP-285/ MPC-200, Sutter Instruments, Novato, CA, USA). Thalamic projections and medial dorsal pallium neurons were visualized with Nikon imaging software (NIS Elements) driving an imaging camera (Hamamatsu ORCA Flash 4.0LT sCMOS, Hamamatsu Photonics, Hamamatsu City, Japan).

To record miniature excitatory postsynaptic currents (mEPSCs), recorded dorsal pallium neurons were confirmed by response to presynaptic stimulation. Slices were then exposed to 250 nM TTX for two minutes. mEPSCs were recorded in 30-second windows of low-noise recordings at -92 mV. Negative -92 mV was selected, as it is near the Cl^-^ reversal potential, which freed our recordings from inhibitory postsynaptic currents.

### Data analysis

Time until energetic failure in extracellular recordings was analyzed by the time elapsed between initial anoxic onset and the final confirmed spontaneous rhythmic burst in both brainstem and spinal cord recordings. Bursts were confirmed by the triangular shape of the integral trace. For analysis on evoked currents, peak amplitude was determined by taking the average of evoked peaks within 60 seconds in their respective time frame, while omitting failed evoked currents. For recovery after high-frequency stimulation, average evoked peak amplitudes were binned in 15-second windows. Amplitudes of evoked currents were detected with the Peak Analysis extension in LabChart (ADInstruments, Dunedin, New Zealand). mEPSCs were analyzed in 30-second timeframes using event detection in Easy Electrophysiology (Easy Electrophysiology, London, UK) with a minimum cut-off of 5 pA. Amplitude values greater than the primary mode of occurrences in each recording were additionally cut off.

### Statistical analysis

For extracellular recordings, time until energetic failure was compared between spinal cords and brainstems before and after hibernation *via* a 2-way ANOVA. Holm-Sidak multiple comparisons tests were then performed to determine changes before and after hibernation, as well as between spinal cord and brainstem network activity. Changes in evoked current from OGD onset was compared after hibernation by running a Mann-Whitney *U* test of the ratio of evoked amplitude in OGD over the baseline evoked amplitude, and compared against an adjusted p-value from a Bonferroni correction. Currents evoked in OGD normalized against their respective normoxic amplitudes were run in one-sample t-tests against the theoretical mean of 1, where no change in evoked amplitude would occur. For mEPSC amplitude and frequency data, changes relative to baseline were compared using Mann-Whitney *U* tests. All statistics were performed using GraphPad Prism (San Diego, CA, USA).

## Results

Previous work has shown the respiratory network output in the American bullfrog brainstem becomes more robust to anoxic exposure after hibernation (Bueschke, Amaral-Silva et al., 2021). To determine if the spinal cord produces a similar response, brainstem and the spinal cord output in response to anoxia was recorded concurrently. Spontaneous rhythmic output was recorded in the isolated brainstem-spinal cord by extracellular recordings from nerve rootlets in the brainstem region that innervate respiratory muscles (trigeminal, vagus, and hypoglossal; Baghdadwala, 2016) and the spinal cord region that outputs rhythmic bursts that are out-of-phase from respiratory activity (SNIII, SNVII-SNIX; Fig 1A-B). Asynchronous activity observed on both nerves suggest that spinal motor activity is independent from the brainstem. We noted occasional large bursts that occurred in-phase between the brainstem and spinal cord. However, the recorded amplitude and burst shape did not match the more consistent rhythmic output of either network.

**Figure 1.**
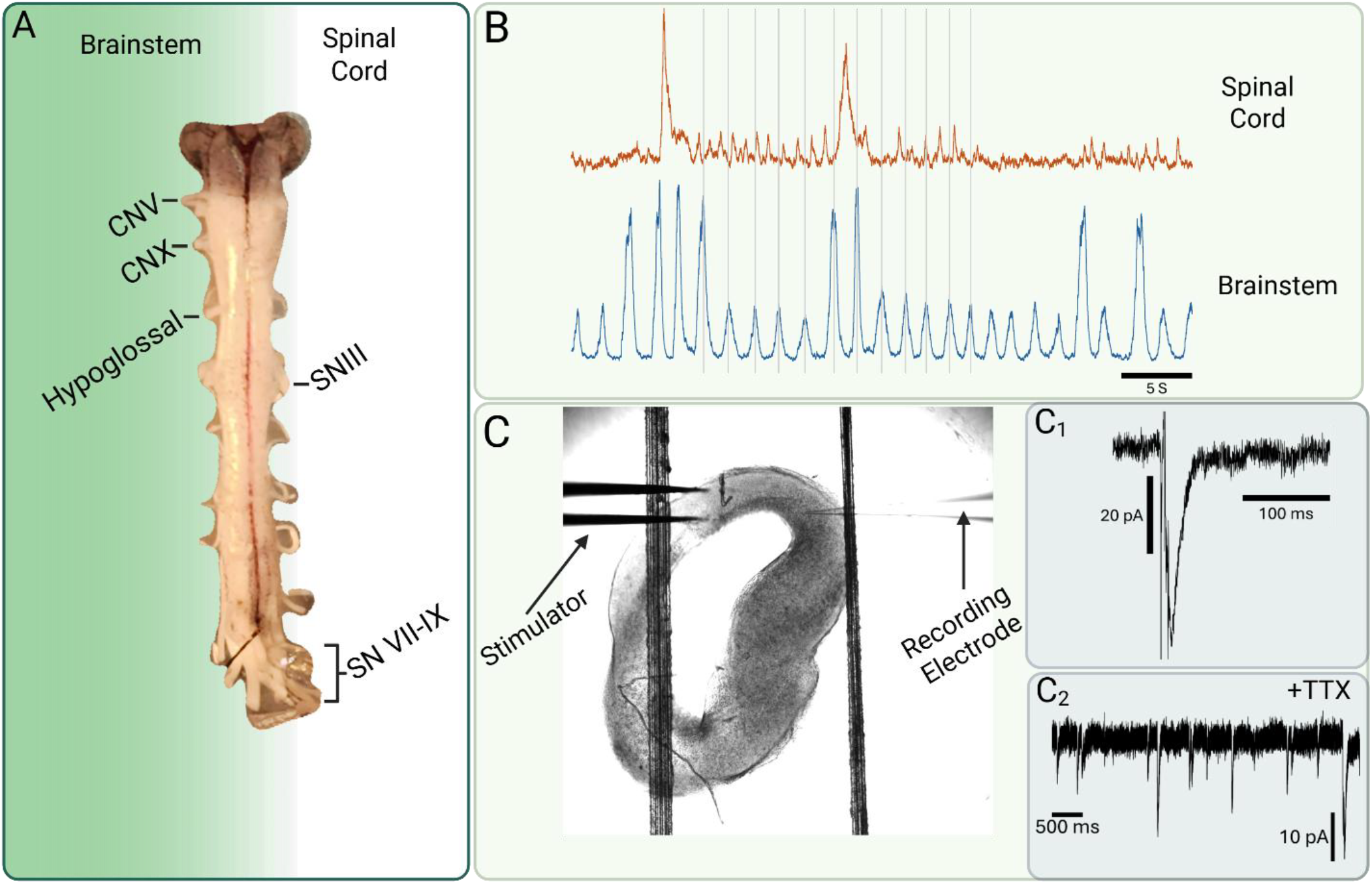
Experimental Approach. (A) Image of an isolated brainstem and spinal cord, ventral side up with nerve rootlets for recordings labeled where respiratory-associated rootlets in the brainstem are shown on the left, and rootlets that output spontaneous spinal cord activity are shown on the right. (B) Representative trace of a simultaneous recording of both spinal cord (top) and brainstem (bottom). Gray lines indicate brainstem bursts to demonstrate that network activity is occurring asynchronously to the spinal cord activity. (C) Image (10X magnification) of a transverse forebrain slice with a stimulator placed over thalamic projections that innervate dorsal pallium neurons, recorded from patch clamp electrodes. (C_1_) Example trace of a stimulator-evoked EPSC in a recorded dorsal pallium neuron. (C_2_) Example trace of an mEPSC recording in the dorsal pallium neuron.

We determined the time until rhythmic failure in anoxia to compare functional response in brainstem and spinal cord activity. We chose to assess this variable, as the time until failure represents the point where the energy available can no longer support neural activity. In un-hibernated control animals, spinal cord function persisted for longer than respiratory activity (Fig 2, p = 0.03; Holm-Sidak’s multiple comparisons test). We found that hibernation improved network function in both the brainstem and the spinal cord (Fig 2; brainstem, p < 0.0001, Holm-Sidak’s multiple comparisons; spinal cord, p < 0.0001, Holm-Sidak’s multiple comparisons), and the difference in anoxic response between the spinal cord and brainstem became more apparent (Fig 2, p = 0.0083; Holm-Sidak’s multiple comparisons). All preparations in controls and hibernators recovered activity following the return of O_2_. Together, these data suggest that the rhythmic network in the spinal cord works independently from the respiratory output observed in the brainstem, but also hibernation induces metabolic changes to increase functional robustness of each network in anoxia.

**Figure 2.**
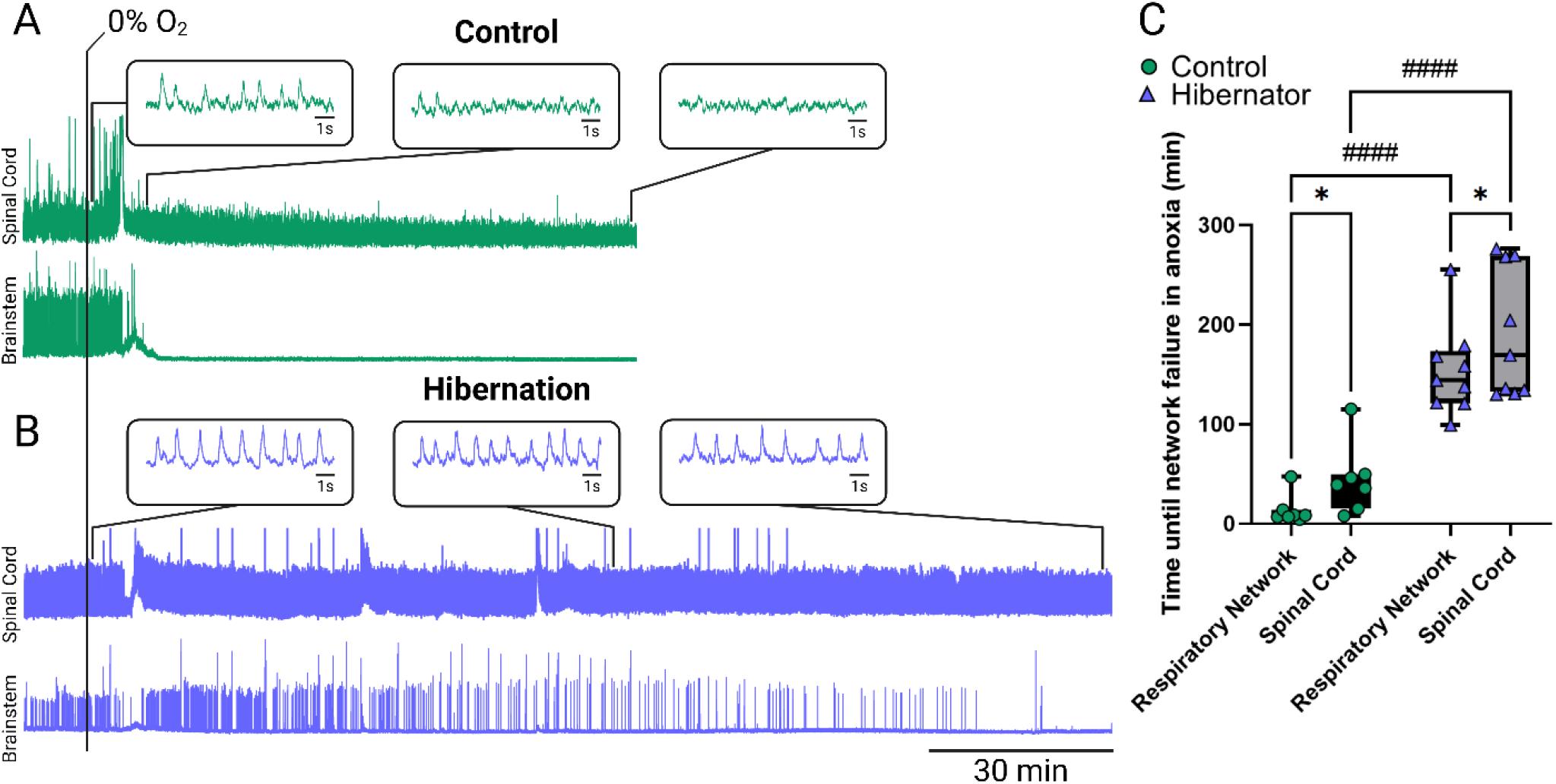
Metabolic plasticity phenotype in the brainstem is present in the spinal cord. (A-B) Representative traces of spontaneous rhythmic output from the brainstem and spinal cord rootlets in the presence of anoxia (0% O_2_) with timing on the same scale between groups. (A) Anoxic response in control conditions shows a rapid cessation of activity in the brainstem. Failure of spinal cord activity was slightly delayed, and was shown as a steady decrease in rhythmic burst amplitude as shown in inlets. (B) Anoxic response post-hibernation shows prolonged brainstem function and spinal cord function that persists for the entire anoxic duration. (C) Time until final duration in anoxia between brainstem and spinal cord activity before and after hibernation. After hibernation, both brainstem and spinal cords persisted longer in anoxia (*n* = 9) compared to pre-hibernated controls (*n* = 7), as shown from results following a Holm-Sidak’s multiple comparisons test. Additionally, spinal cord activity persisted longer in anoxia than brainstems before and after hibernation.

To determine if these responses were unique to the brainstem and spinal cord, we addressed neuronal function of neurons from medial dorsal pallium neurons in the forebrain. We focused on synaptic function since previous studies demonstrated that membrane potential and firing capacity in pallium neurons is unaffected in anoxia in both un-hibernated controls and hibernators (Amaral-Silva & Santin, 2023). Additionally, synapses were the primary limit to circuit function in brainstem motor networks in bullfrogs. To achieve this, we recorded postsynaptic currents that were evoked in response to presynaptic stimulation of thalamic projections that provide input to the dorsal pallium (Roth et al., 2003, 2007). Amplitudes of maximal evoked currents stimulated at 0.5 Hz were recorded in normoxic aCSF and compared after 15 minutes in oxygen and glucose deprivation (OGD). While controls and hibernators had no difference in baseline amplitude in normoxia (Fig 3C; p = 0.3604; Holm-Sidak’s multiple comparisons test), both control and hibernator groups had a significant change in amplitude in OGD (Fig 3B; Control, t_(29)_ = 11.79; p < 0.0001; one-sample *t* test; Hibernator, t_(30)_ = 3.512; p = 0.0014; one-sample *t* test) where OGD resulted in evoked synaptic currents that were an average of 57% the baseline amplitude in controls and 81% after hibernation. OGD-induced decreases in amplitude being present in both control and hibernated animals were also shown when comparing recorded evoked EPSCs, given a significant effect on condition (F_(1,59)_ = 27.43; p < 0.0001; two-way repeated measures ANOVA) and significant changes between conditions in both groups (Fig 3C; Controls, p < 0.0001; Hibernators, p = 0.0174; Holm-Sidak’s multiple comparisons test). Despite both groups decreasing synaptic currents to some extent, when assessing the change during OGD, hibernators better maintained evoked synaptic currents compared to controls, decreasing less (p = 0.0016; Mann-Whitney *U* test with Bonferroni corrected p-value). To further assess how synaptic transmission was affected, we made comparisons of miniature excitatory postsynaptic current (mEPSC) amplitude and frequency changes in OGD before and after winter. Neither mEPSC amplitude (Fig 4A; p = 0.1802, Mann-Whitney *U* test), nor frequency (Fig 4B; p = 0.5824, Mann-Whitney *U* test) showed differences in response to OGD in dorsal pallium neurons.

**Figure 3.**
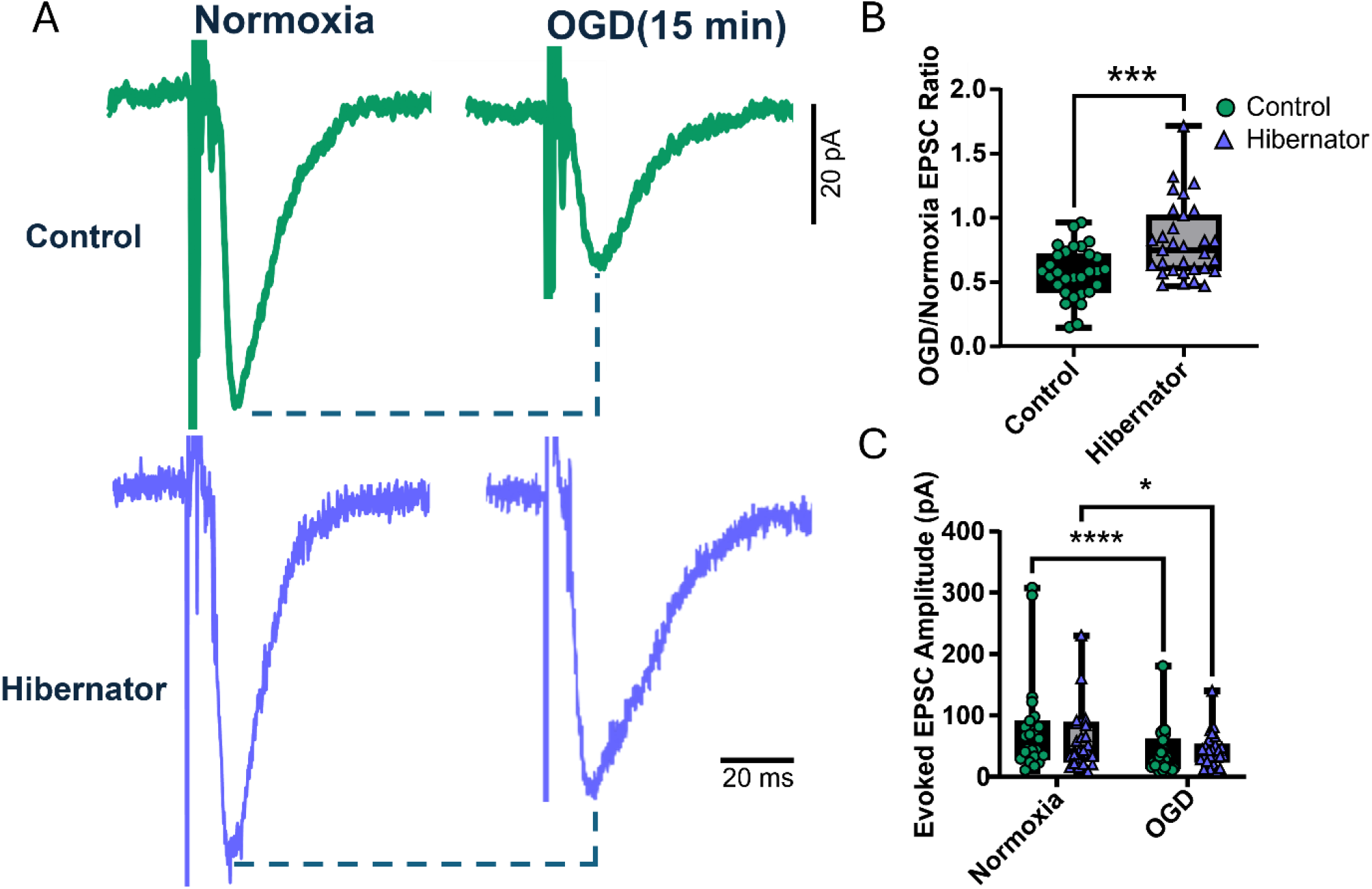
Hibernation decreases sensitivity to oxygen and glucose deprivation in dorsal pallium synapses. OGD onset results in attenuated evoked EPSC amplitudes that are less severe after hibernation. (A) Example traces of individual evoked currents comparing amplitudes before and after 15 minutes of OGD exposure in control (top) and after hibernation (bottom). (B) Evoked EPSC amplitude in OGD relative to baseline in controls (*n* = 30 neurons, *N* = 12 animals) and hibernators (*n* = 31 neurons, *N* = 15 animals). Significance in amplitude changes was shown with a Mann-Whitney *U* test. (C) Recorded evoked EPSC amplitudes to compare differences in absolute amplitudes between groups. Holm-Sidak’s multiple comparisons show significance in both controls and hibernators.

**Figure 4.**
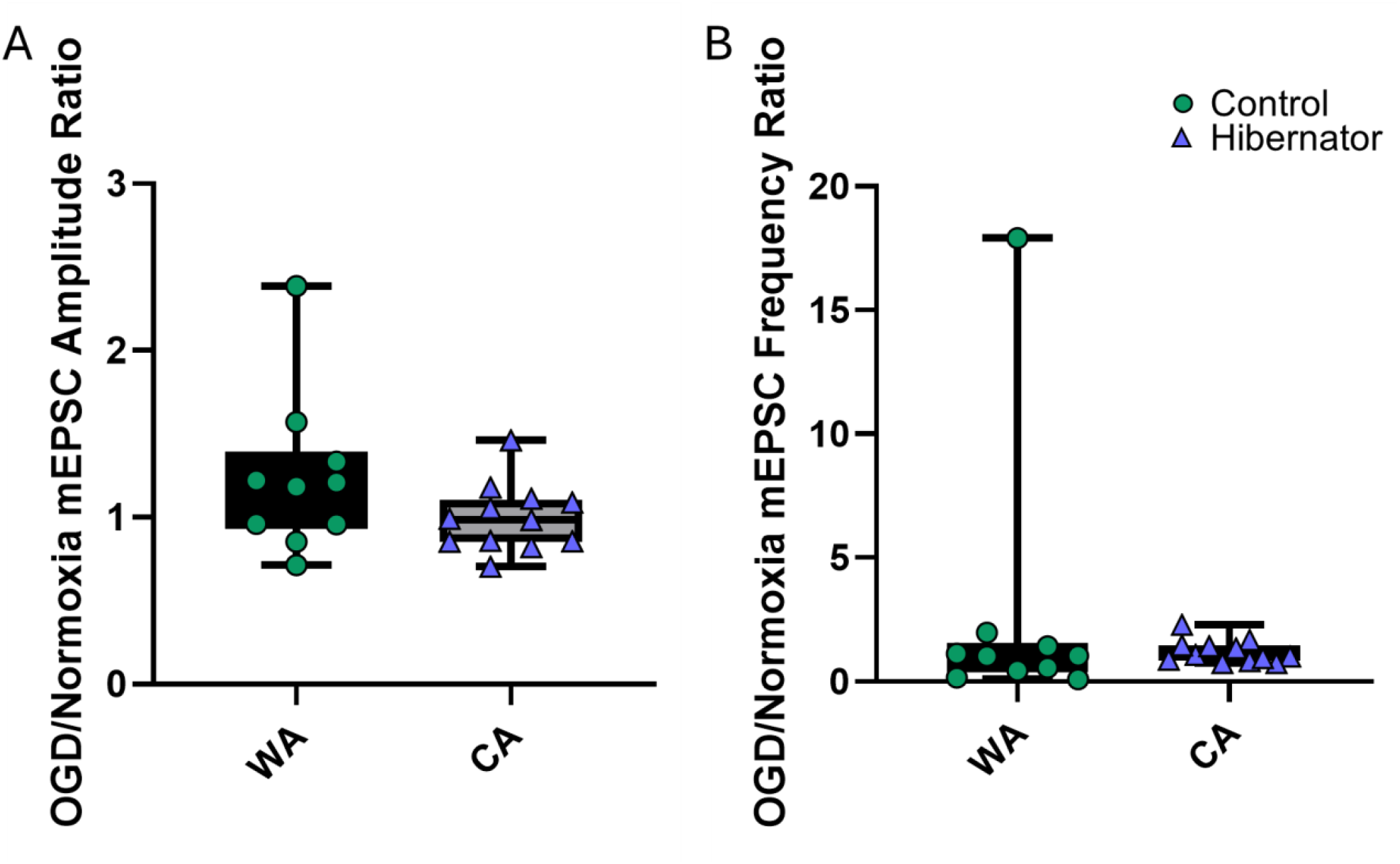
Hibernation does not alter mEPSC amplitude or frequency. (A) Change in mEPSC amplitxsude in 15 minutes of OGD relative to normoxic conditions. Mann-Whitney *U* results showing no difference in mEPSC amplitude suggest that OGD has no influence on changes in postsynaptic response to excitatory quanta. (B) Difference in mEPSC frequency in OGD relative to normoxic conditions. Mann-Whitney *U* test results showing no changes in altered frequency after hibernation suggest no change in presynaptic release sites.

## Discussion

Given the high energy demands of active neural circuits, function of the CNS rapidly fails during severe energy deficit (Somjen, 2001). Although amphibians are not generally considered to be strongly hypoxia-tolerant, we have previously demonstrated that the American bullfrog can transform rhythmic brainstem circuits to maintain activity during prolonged exposure to anoxia and ischemia after hibernation (Amaral-Silva & Santin, 2023; Bueschke, Amaral-Silva et al., 2021). Here, we investigated how different regions of the CNS of the frog respond to energy stress to determine if the respiratory motor network is unique in the metabolic plasticity phenotype to improve tolerance to stress. Our results suggest that the improvement in neuronal function during severe energy deficits after hibernation occurs throughout the CNS.

### Comparison of control response to energy stress across networks

Prior to hibernation in controls, we observed differences in responses between the brainstem and spinal cord, consistent with regional differences in sensitivity to hypoxia. Spontaneous rhythmic activity in the spinal cord persists longer in anoxia in control frogs compared to activity from the respiratory motor network in the brainstem (Fig 2). This suggests that the spinal cord may be more effective at maintaining function in glycolysis through either through higher glycolytic rates or decreased energy demands based on differences in network architecture and design (Quintela-López et al., 2022). Additionally, while synapses in the dorsal pallium indeed have a notable decrease in evoked EPSC amplitude in OGD in control frogs, the response is much less dramatic when compared to brainstem motor neurons from the vagal motor pool in the same species. Indeed, anoxia rapidly attenuates the entire evoked EPSCs within 10 minutes (Amaral-Silva & Santin, 2023). In contrast, evoked EPSCs in the dorsal pallium decreased to an average of 57% of the baseline current after 15 minutes without oxygen or glucose (Fig 3). While there are differences in the way these experiments were performed (e.g., direct stimulation of presynaptic terminals near the cell body vs. more distant axons), work in mammals demonstrated that neurons in the neocortex have a much more delayed response to anoxic depolarization, a staple of neuronal failure in energy stress, compared to hypoglossal motor neurons in the brainstem of rats (O’Reilly et al., 1995). Therefore, at baseline, we observed differences in responses to energy stress in different parts of the central nervous system.

### Comparison of improvements following hibernation

Despite differences in the responses of each network to energy stress in control animals, these three networks have much more prominent similarities when considering their improvements after hibernation. Indeed, we built upon our past work in the brainstem and demonstrated that spinal and forebrain neurons both improve tolerance to severe energy stress. While the brainstem and spinal cord improved, we observed quantitative differences in these responses. After hibernation, the respiratory motor network maintained function in anoxia for approximately 11-fold longer than controls, while the average improvement was approximately 5-fold in the spinal cord (Fig 2). However, this is likely to be an underestimate as 7 out of 9 spinal cords were still active when we terminated the experiments, well after respiratory activity failed. In brainstem motoneurons shown in Amaral-Silva and Santin (2023), hibernation led to maintained synaptic function in anoxia with no difference compared to baseline for up to 40 minutes, with one exceptional recording even lasting 2 hours with maintained synaptic transmission (Amaral-Silva & Santin, 2023). Notwithstanding the differences between anoxia and OGD and differences in the specific preparations used in these studies, the picture seems to be different in the dorsal pallium, where evoked synaptic transmission significantly decreased by ∼20% after 15 minutes, despite performing better than control animals at the same time point (Fig 3B). Therefore, while each part of the nervous system appears to improve its tolerance to energy stress, they differ in the degree to which this occurs. The mechanisms that give rise to these differences are not yet clear but may involve the extent to which glycolysis can take over as a lone fuel source without aerobic respiration. Previous work suggests that this is controlled by a transcriptional program that coordinates multiple metabolic responses including glycogen metabolism, glucose transport, and glycolysis (Hu & Santin, 2022). Therefore, differences in the capacity for the hibernation environment to alter metabolic gene expression may drive some of the differences across brain regions.

### Perspectives and significance

Breathing is essential for metabolic homeostasis and gas exchange in air-breathing animals. However, frogs in the hibernation environment perform only cutaneous gas exchange, as they are submerged in cold water (Santin & Hartzler, 2017; Tattersall & Ultsch, 2008; Ultsch et al., 2004). In this environment, blood O_2_ is low, which poses a problem for ramping up neural activity that is needed to restore metabolic homeostasis as the temperature warms during emergence. Therefore, initial work focused on the brainstem respiratory network, as this circuit along with those controlling cardiovascular function, are most vital for restoring metabolic homeostasis. Given the striking differences in the capacity to function during anoxia and ischemia after hibernation, this led us to hypothesize that the brainstem respiratory network may be “prioritized” in becoming resistant to energy stress (i.e., the respiratory neurons can turn on under hypoxia to restore breathing, which achieves metabolic homeostasis of the rest of the body and brain as the animal emergences). However, our results here suggest that improvements in neural performance are more widespread, extending to rhythmic spinal motor circuits and areas of the forebrain controlling associative learning. While we did not exhaustively sample a large number of brain and spinal cord regions, these results nevertheless suggest that the forebrain, brainstem, and spinal cord have the capacity to improve their tolerance to energy stress. Therefore, metabolic plasticity is likely to aid in other behaviors that must occur on the background of hypoxia during emergence beyond lung ventilation, such as locomotion or various behavioral tasks. More broadly, we assert that understanding the mechanisms that explain quantitative differences observed across regions may lead to new mechanistic insights. Is each region and cell type using similar mechanisms, or are there multiple ways to become hypoxia tolerant even within the same individual? By investigating these processes and comparing results across a variety of interesting hypoxia-tolerant species, we will gain greater insights into new ways that the brain may alter its metabolic requirements.

## Acknowledgment

We would like to thank the National Institutes of Health (#R01NS114514 and R15NS112920) for funding.

## References

Adams, S., Zubov, T., Bueschke, N., & Santin, J. M. (2021). Neuromodulation or energy failure? Metabolic limitations silence network output in the hypoxic amphibian brainstem. American Journal of Physiology-Regulatory, Integrative and Comparative Physiology, 320(2), R105–R116. 10.1152/ajpregu.00209.2020

Amaral-Silva, L., & Santin, J. (2024). Neural Processing without O _2_ and Glucose Delivery: Lessons from the Pond to the Clinic. Physiology, 39(6), 000–000. 10.1152/physiol.00030.2023

Amaral-Silva, L., & Santin, J. M. (2023). Synaptic modifications transform neural networks to function without oxygen. BMC Biology, 21(1), 54. 10.1186/s12915-023-01518-0

Ashton, D., Van Reempts, J., Haseldonckx, M., & Willems, R. (1989). Dorsal-ventral gradient in vulnerability of CA1 hippocampus to ischemia: A combined histological and electrophysiological study. Brain Research, 487(2), 368–372. 10.1016/0006-8993(89)90842-1

Baghdadwala, M. I. (2016). Diving into the mammalian swamp of respiratory rhythm generation with the bullfrog. Respiratory Physiology.

Brisson, C. D., Hsieh, Y.-T., Kim, D., Jin, A. Y., & Andrew, R. D. (2014). Brainstem Neurons Survive the Identical Ischemic Stress That Kills Higher Neurons: Insight to the Persistent Vegetative State. PLoS ONE, 9(5), e96585. 10.1371/journal.pone.0096585

Buck, L. T., & Pamenter, M. E. (2018). The hypoxia-tolerant vertebrate brain: Arresting synaptic activity. Comparative Biochemistry and Physiology Part B: Biochemistry and Molecular Biology, 224, 61–70. 10.1016/j.cbpb.2017.11.015

Bueschke, N., Amaral-Silva, L. do, Adams, S., & Santin, J. M. (2021). Transforming a neural circuit to function without oxygen and glucose delivery. Current Biology, 31(24), R1564–R1565. 10.1016/j.cub.2021.11.003

Bueschke, N., Amaral-Silva, L., Hu, M., Alvarez, A., & Santin, J. M. (2024). Plasticity in the functional properties of NMDA receptors improves network stability during severe energy stress. The Journal of Neuroscience, e0502232024. 10.1523/JNEUROSCI.0502-23.2024

Fernandes, J. A., Lutz, P. L., Tannenbaum, A., Todorov, A. T., Liebovitch, L., & Vertes, R. (1997). Electroencephalogram activity in the anoxic turtle brain. American Journal of Physiology-Regulatory, Integrative and Comparative Physiology, 273(3), R911–R919. 10.1152/ajpregu.1997.273.3.R911

Hofmeijer, J., Franssen, H., Van Schelven, L. J., & Van Putten, M. J. A. M. (2013). Why Are Sensory Axons More Vulnerable for Ischemia than Motor Axons? PLoS ONE, 8(6), e67113. 10.1371/journal.pone.0067113

Howell, B., Baumgardner, F., Bondi, K., & Rahn, H. (1970). Acid-base balance in cold-blooded vertebrates as a function of body temperature. American Journal of Physiology-Legacy Content, 218(2), 600–606. 10.1152/ajplegacy.1970.218.2.600

Hu, M., & Santin, J. M. (2022). Transformation to ischaemia tolerance of frog brain function corresponds to dynamic changes in mRNA co-expression across metabolic pathways. Proceedings of the Royal Society B: Biological Sciences, 289(1979), 20221131. 10.1098/rspb.2022.1131

Hurley, E. P., Mukherjee, B., Fang, L., Barnes, J. R., Nafar, F., Hirasawa, M., & Parsons, M. P. (2022). Atypical NMDA Receptors Limit Synaptic Plasticity in the Adult Ventral Hippocampus [Preprint]. Neuroscience. 10.1101/2022.10.05.510966

Jackson, D. C., & Ultsch, G. R. (2010). Physiology of hibernation under the ice by turtles and frogs. Journal of Experimental Zoology Part A: Ecological Genetics and Physiology, 313A(6), 311–327. 10.1002/jez.603

Jinka, T. R., Rasley, B. T., & Drew, K. L. (2012). Inhibition of NMDA‐type glutamate receptors induces arousal from torpor in hibernating arctic ground squirrels (Urocitellus parryii). Journal of Neurochemistry, 122(5), 934–940. 10.1111/j.1471-4159.2012.07832.x

Larson, J., Drew, K. L., Folkow, L. P., Milton, S. L., & Park, T. J. (2014). No oxygen? No problem! Intrinsic brain tolerance to hypoxia in vertebrates. Journal of Experimental Biology, 217(7), 1024–1039. 10.1242/jeb.085381

O’Reilly, J. P., Jiang, C., & Haddad, G. G. (1995). Major differences in response to graded hypoxia between hypoglossal and neocortical neurons. Brain Research, 683(2), 179–186. 10.1016/0006-8993(95)00373-X

Quintela-López, T., Shiina, H., & Attwell, D. (2022). Neuronal energy use and brain evolution. Current Biology, 32(12), R650–R655. 10.1016/j.cub.2022.02.005

Rauscent, A., Le Ray, D., Cabirol-Pol, M.-J., Sillar, K. T., Simmers, J., & Combes, D. (2006). Development and neuromodulation of spinal locomotor networks in the metamorphosing frog. Journal of Physiology-Paris, 100(5–6), 317–327. 10.1016/j.jphysparis.2007.05.009

Roth, G., Grunwald, W., & Dicke, U. (2003). Morphology, axonal projection pattern, and responses to optic nerve stimulation of thalamic neurons in the fire-bellied toad Bombina orientalis. The Journal of Comparative Neurology, 461(1), 91–110. 10.1002/cne.10670

Roth, G., Laberge, F., Mühlenbrock-Lenter, S., & Grunwald, W. (2007). Organization of the pallium in the fire-bellied toad Bombina orientalis. I: Morphology and axonal projection pattern of neurons revealed by intracellular biocytin labeling. The Journal of Comparative Neurology, 501(3), 443–464. 10.1002/cne.21255

Santin, J. M., & Hartzler, L. K. (2017). Activation of respiratory muscles does not occur during cold-submergence in bullfrogs, Lithobates catesbeianus. Journal of Experimental Biology, jeb.153544. 10.1242/jeb.153544

Somjen, G. G. (2001). Mechanisms of Spreading Depression and Hypoxic Spreading Depression-Like Depolarization. Physiological Reviews, 81(3), 1065–1096. 10.1152/physrev.2001.81.3.1065

Sotelo, M. I., Daneri, M. F., Bingman, V. P., & Muzio, R. N. (2024). Amphibian spatial cognition, medial pallium and other supporting telencephalic structures. Neuroscience & Biobehavioral Reviews, 163, 105739. 10.1016/j.neubiorev.2024.105739

Sutherland, R. M., & Nunnemacher, R. F. (1974). Fibers in the ventral spinal nerves of the frog. Journal of Comparative Neurology, 156(1), 39–47. 10.1002/cne.901560105

Tattersall, G. J., & Ultsch, G. R. (2008). Physiological Ecology of Aquatic Overwintering in Ranid Frogs. Biological Reviews, 83(2), 119–140. 10.1111/j.1469-185X.2008.00035.x

Ultsch, G. R., Reese, S. A., & Stewart, E. R. (2004). Physiology of hibernation inRana pipiens: Metabolic rate, critical oxygen tension, and the effects of hypoxia on several plasma variables. Journal of Experimental Zoology, 301A(2), 169–176. 10.1002/jez.a.20014

Woych, J., Ortega Gurrola, A., Deryckere, A., Jaeger, E. C. B., Gumnit, E., Merello, G., Gu, J., Joven Araus, A., Leigh, N. D., Yun, M., Simon, A., & Tosches, M. A. (2022). Cell-type profiling in salamanders identifies innovations in vertebrate forebrain evolution. Science, 377(6610), eabp9186. 10.1126/science.abp9186

Zhao, H. W., Ross, A. P., Christian, S. L., Buchholz, J. N., & Drew, K. L. (2006). Decreased NR1 Phosphorylation and Decreased NMDAR Function in Hibernating Arctic Ground Squirrels. Journal of Neuroscience Research, 84(2), 291–298. 10.1002/jnr.20893

